# Mapping spatial cell-cell communication programs by tailoring chains of cells for transformer neural networks

**DOI:** 10.64898/2026.03.18.712664

**Authors:** Niklas Brunn, Laia C. Guitart, Kiana Farhadyar, Camila L. Fullio, Jakob Kailer, Tanja Vogel, Maren Hackenberg, Harald Binder

## Abstract

Recent advances in spatial transcriptomics and computational modeling enable the study of cellular interactions *in situ*. However, existing methods quantify ligand–receptor activity pairwise or between predefined cell groups, yielding overlapping signals and limited ability to summarize concurrent interactions into programs while localizing communication hotspots. We introduce scCChain, a transformer-based framework that integrates ligand–receptor activity into spatially resolved communication programs and localizes hotspots at spot and single-cell resolution. scCChain first derives candidate programs using structured dimensionality reduction. Subsequently, it samples program-specific communication chains by linking transcriptionally similar sender cells to candidate receivers via weighted random walks on a distance-informed cell graph, borrowing signal from similar neighbors. Transformer-based modeling then scores chains to prioritize communication programs and pinpoint hotspots across the tissue. Applied to human breast cancer spatial transcriptomics data at spot and single-cell resolution, scCChain supports both exploratory communication program discovery and targeted analysis of user-specified ligand–receptor pairs. In spot-level data, it prioritizes a tumor-associated program enriched for pro-angiogenic signaling that localizes to invasive regions. In imaging-based data, it highlights CXCL12–CXCR4 communication hotspots at cellular resolution. Here, we demonstrate that chain-based transformer modeling enables interpretable discovery and mapping of biological meaningful spatial communication programs within complex tissues.

## Introduction

Cell-cell communication (CCC) is fundamental to a range of processes, including development [1, 2], tissue homeostasis [3], and disease [4, 5, 6]. Mechanistically, one common principle is ligand–receptor signaling via secreted factors, in which sender cells release ligands that bind to cognate receptors on receiver cells to trigger downstream gene-regulatory responses [7, 8]. The spatial context shapes these signals [7, 9], and recent spatial transcriptomics assays map gene expression *in situ*, ranging from spot-level to subcellular resolution [10, 11, 12]. Combined with curated ligand–receptor resources and pioneering computational approaches [13, 14, 15, 16], this has enabled modeling of spatially organized CCC [17, 18, 19, 20, 21].

However, CCC is rarely mediated by isolated ligand–receptor pairs. Instead, multiple pairs organize into communication programs (CPs) that work in concert [7, 22]. Examples include pathway-level crosstalk, where multiple ligands and receptors operate within the same or convergent pathways [7, 23], and relay networks, in which one interaction triggers the next [21]. Modeling CPs rather than isolated pairs therefore better reflects tissue biology [7, 23]. Although some computational approaches have already touched on CPs [22, 24], they are not tailored to spatial transcriptomics and limited to modeling CCC in cell pairs, thereby neglecting the complexity of signaling information from the microenvironment. To this end, we introduce scCChain (single-cell communication chains), a chain-based framework that enables the use of the powerful transformer neural network architecture [25] to identify spatial communication programs at single-cell and spot resolution. scCChain reframes CCC as a sequence-modeling problem, in which weighted random walks on a distance-aware cell graph assemble chains of cells. Two complementary edge types drive the graph: (i) similarity edges that connect spatially proximate, transcriptionally similar cells (guilty-by-association) and (ii) ligand–receptor interaction edges linking putative sender and receiver cells based on coexpression of ligand–receptor pairs from a curated database. To summarize ligand–receptor activity into CPs, we apply structured dimensionality reduction [26], recently adapted to CCC data [27]. This yields CPs characterized by sparse loadings over ligand–receptor pairs, which guide communication chain construction. Further assuming that a receiver cell’s transcriptome reflects downstream responses shaped by the gene expression states of its sender cells, we expand each cell’s feature representation within a chain to include genes beyond ligand–receptor databases, enabling transformer-based prioritization of CPs.

Currently, most computational CCC approaches model cellular interactions either as individual cell pairs [18, 22], or via spatial graphs that expand neighborhoods to capture a multi-hop context [17, 19, 21]. Pairwise modeling is brittle under the noise typical for spatial transcriptomics data [28, 29] and neglects spatial environmental dependencies [17]. In contrast, graph models expand neighborhoods with radius or depth (multi-hop), but even modest radius increases rapidly include large node sets, raising computational costs on large graphs [30, 31]. A complementary, yet unexplored alternative is to cast CCC as a sequence-modeling problem. Chains of cells can traverse a tissue over variable distances, while keeping the effective number of cells per chain small and still being able to integrate information from the microenvironment. The chain construction mirrors how a human expert might trace CCC through tissue by jointly integrating spatial, transcriptional, and directed signaling information. Each chain starts from a candidate sender cell for a specific communication mechanism and traverses nearby transcriptionally similar cells to borrow information under sparse and noisy spatial transcriptomics measurements. Along the way, it searches for a candidate receiver cell by tracking expression of mechanism-relevant genes along the chain.

Despite the rich information in whole-transcriptome single-cell and spatial assays, current technologies cannot observe communication events directly. As a result, a widely accepted ground truth for CCC is missing, which prevents fully supervised training and complicates the evaluation of model-derived patterns [7, 16]. In practice, most studies rely on indirect evidence such as agreement across tools, spatial proximity, protein measurements, incorporation of downstream signaling information, and on targeted perturbations or other resource-intensive experimental follow-ups [16, 22, 32]. Because computational methods for modeling CCC depend on curated ligand–receptor databases, analyses are restricted to active genes represented in those resources and may omit relevant signals outside the catalogued set. At the same time, a receiver cell is influenced by its local niche, and the expression of signaling-responsive genes is conditionally dependent on the states of nearby sender cells [17]. Recent work further indicates that leveraging cross-cellular dependency structure in gene expression can be sufficient to recover tissue organization from dissociated profiles alone [33]. These kinds of patterns, together with cellular interactions, can be reflected in a sequence prediction task, where the gene expression of a cell is predicted by the gene expression of prior cells in a chain. We implement this dependency using transformer neural networks [25, 34]. Through multi-head self-attention, a transformer model can selectively weight informative cells and gene signals along the chain and capture coordinated whole-transcriptome patterns across chains in receiver cells that are consistent with the expression states of their putative senders. From this chain perspective, the performance of a fitted prediction model provides an interpretable criterion for communication plausibility, where lower prediction errors indicate higher plausibility. We therefore use this signal to prioritize candidate CCCs for CPs or individual ligand–receptor pairs.

For illustrating this chain perspective and the specific implementation via transformers, we apply the proposed scCChain approach to matched spot-level and single-cell–resolution spatial transcriptomics datasets from human breast cancer tissue. In Visium CytAssist data, scCChain prioritizes a communication program enriched for VEGF-, midkine- and WNT-related ligand–receptor pairs that is predominantly localized to invasive tumor regions, consistent with a pro-angiogenic, tumor-associated signaling module. In corresponding Xenium In Situ data, we use a targeted analysis of CXCL12–CXCR4 signaling to characterize cell-type-specific sender-receiver relationships and sender-receiver distances. Together, these examples illustrate that scCChain aggregates ligand–receptor pairs into spatially localized communication programs represented by chains, and that transformer-based prioritization highlights plausible communication hotspots, their dominant sender-receiver cell types, and characteristic spatial ranges.

## Results

### Spatial localization and prioritization of cell-cell communication programs with scCChain

In tissues, local microenvironments shape cellular behavior, and cells interact through coordinated communication programs (CPs) [22, 24] of ligand–receptor pairs [7, 23]. We developed scCChain to localize and prioritize CP activity within tissue samples.

The framework operates on spatial transcriptomics data generated using diverse technologies [10, 11, 12]. It requires gene expression counts of either single cells (for example, after cell segmentation) or spatial regions (spots or bins), together with their center coordinates (Fig. 1a). By default, we use CellChatDB [14] as the reference catalogue of ligand–receptor pairs, but other curated or custom databases are also supported. The modeling framework does not require prior cell type annotations, although such labels can aid visualization and interpretation of the results.

**Figure 1:**
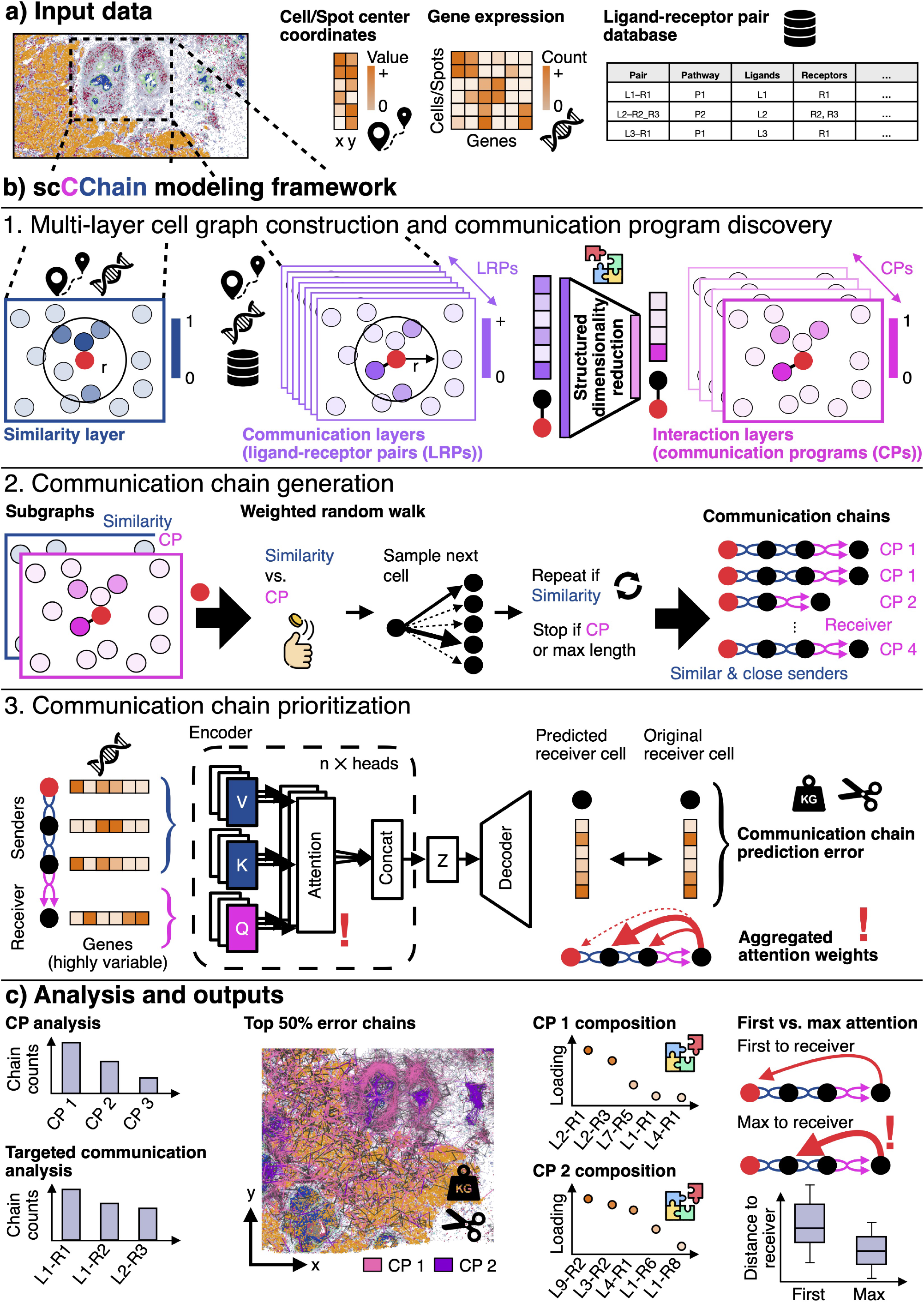
scCChain overview. **a)** Inputs to scCChain include spatial coordinates of cells or spots, gene expression counts, and a curated ligand receptor database. **b)** Modeling framework. 1. A distance-aware cell graph is constructed with a transcriptional similarity layer and directed communication layers that encode distance-weighted ligand and receptor coexpression between putative sender receiver pairs. Structured dimensionality reduction across ligand–receptor pairs groups concordant communication patterns into communication programs with sparse positive ligand–receptor pair loadings. 2. For each communication program, communication chains are sampled by weighted random walks on the corresponding subgraph, alternating between similarity and communication edges until a communication step occurs or a maximum chain length is reached. 3. Chains are prioritized by fitting a transformer model that predicts receiver gene expression from sender cells along the chain. The receiver is used as the query and sender cells are used as keys and values. Chain-level prediction error ranks candidate communication events. Attention weights highlight influential senders. **c)** Outputs include communication program discovery or targeted ligand–receptor pair analysis by omitting the program discovery step, spatial mapping of prioritized chains to localize communication hotspots with line width and opacity scaled inversely with prediction error, communication program compositions from ligand–receptor pair loadings, and attention-based sender summaries, including first sender-to-receiver and max-attention sender-to-receiver distances.

As a basis, we construct a distance-aware cell graph with two kinds of edge layers, the similarity layer and the communication layers (Fig. 1b, 1). Similarity edges are weighted by transcriptional similarity, smoothed with a truncated distance kernel in a way that spatially proximate cells are favored during sampling. Communication layers encode directed communication potential for all ligand–receptor pairs from the database, again smoothed by the distance kernel (Fig. 1b, 1). Communication potential is modeled from coexpression of cognate ligands and receptors in sender-receiver pairs.

To identify CPs, we apply structured dimensionality reduction [26, 27] to the communication layer edge weights across the axis of ligand–receptor pairs for spatially constrained cell pairs. This yields a low-dimensional representation in which each latent component defines a CP, characterized by a sparse, positively weighted set of ligand–receptor pairs. Thus, for each cell pair, the corresponding CP value can be interpreted as an aggregated communication score across these ligand–receptor pairs.

The second part of the modeling framework samples communication chains using weighted random walks on the cell graph (Fig. 1b, 2). For each CP-specific communication layer, we build a subgraph comprising that layer together with the similarity layer, and generate chains from sender cells with high outgoing signaling potential. A coin-flip mechanism alternates moves between similarity and communication edges until a communication edge is selected or a maximum chain length is reached, allowing chains to traverse local neighborhoods while remaining concise. The random walk is designed to enrich the chain with cells that are transcriptionally similar to the candidate sender cell, allowing for borrowing of information according to a guilty-by-association strategy. Under this strategy, a cell similar to the candidate sender cell might either be the actual sender cell, with the pattern distorted by noise, or might just be useful for seeing more clearly the pattern around the candidate sender cell. Due to the random walk, a specific cell might be part of several chains. In each of these chains, the cell might become the candidate sender cell, based on its communication potential, i.e. our approach does not require to designate candidate sender cells beforehand. Notably, similarity edges can route chains through cells with low or no ligand expression that would be discarded by pairwise-communication models but may still carry relevant transcriptional information. The resulting enriched chains represent candidate communication events for each CP. To represent each cell along a chain, we use not only genes covered by the ligand–receptor pair database but also a broader set of highly variable genes.

Subsequently, filtering identifies chains and their components that are likely to reflect active communication (Fig. 1b, 3). To this end, we formulate CCC potential across chains as a prediction task, where the objective is to predict the receiver cell’s gene expression from the gene activity of the sender cells within each chain. Intuitively, stronger cellular crosstalk yields stronger cross-gene dependencies between senders and receiver, improving prediction of the receiver’s gene expression. We therefore prioritize candidate CCCs based on chain-wise prediction errors. The chain format enables transformer-based sequence modeling [25], where tokens correspond to cells in the chain and their feature vectors to their gene expression profile. In addition, the attention mechanism underlying transformer-based model architectures enables learning attention weights to quantify the contribution of each sender cell in a chain for predicting the matched receiver cell as a basis for subsequent interpretation.

scCChain provides several outputs that facilitate spatial CCC analysis (Fig. 1c). Although we have specifically designed the framework for analyzing CPs, users can also omit the CP discovery step and instead choose to run the analysis for a preferred set of ligand–receptor pairs. In that case, chains are generated for all feasible ligand–receptor pairs prior to communication prioritization (Fig. 1c). The prioritized communication chains can be mapped to the tissue to localize CCC activity hotspots for different CPs (Fig. 1c). Specifically, chains with a high prediction error can be filtered, and the edge width and transparency for the remaining chains can be weighted inversely by the error values to visually highlight regions prioritized by the transformer model. Additionally, the composition of CPs in different ligand–receptor pairs can be examined by the corresponding loadings derived from the CP dimensionality reduction model (Fig. 1c). The normalized loading scores indicate the contribution strength to the communication patterns captured by different CPs. Further, the attention weights of the fitted transformer model pinpoint influential sender cells for predicting the matched receiver cells (Fig. 1c). The most attended cells can be further inspected for attributes such as the distance to the corresponding receiver cell, which can be contrasted to the first sender cells in the chains (Fig. 1c).

Together, these steps recast spatial CCC as a sequence-modeling problem. This in turn enables transformers to prioritize spatially localized CPs, and highlight influential sender cells. In contrast, many computational approaches act directly on node-centric graphs to model CCC in the tissue [17, 18]. However, acting on node-centric graphs requires setting a fixed radius for which transcripts from other nodes are taken into account, or fixing the number of nodes directly. Increasing the radius leads to an exponential increase of nodes in range (Fig. S1 a), which becomes crucial for modeling single-cell level data. On the other side, relying on a fixed number of neighboring nodes can cause other unwanted effects, since node density is regional and neighbor nodes for different center nodes can have varying ranges (Fig. S1 b). Distance-aware chains of cells, defined as directed paths on the underlying cell graph, circumvent the limitations of fixed, radius- or degree-based neighborhood definitions by re-centering the node-centric view at each sampling step of a random walk (Fig. S1 c). As a consequence, both the number of nodes assembled within each chain and the maximal spatial range a chain can traverse through tissue are controlled by the neighborhood radius, the maximum number of transitions per chain, and the coin flip success probability for selecting the communication layer during node sampling (Fig. S1 d,e). In particular, chains can span longer tissue ranges while keeping the number of collected nodes small. The chain structure is particularly favorable for flexible modeling with artificial neural network architectures. Unlike dense spatial graphs, whose memory and computational modeling demands scale exponentially with the number of nodes, communication chains scale linearly with chain length, making them computationally tractable for transformer architectures [25] that rely on the scaled dot-product attention mechanism. Moreover, chains are ordered inherently, which enables transformer-based modeling, as these are designed to learn dependencies across sequences. We therefore design chains to encode directed communication paths from senders to receivers, enabling the model to attend selectively to influential sender cells.

### scCChain reveals tumor-related communication program hotspots in human breast cancer

To demonstrate how scCChain localizes and prioritizes spatial CPs, we applied it to spot-level Visium CytAssist spatial transcriptomics data from a human breast cancer specimen. The data were initially generated to investigate tumor heterogeneity [10] (Fig. 2a). The preprocessed data consist of 4,982 hexagon-shaped spots arranged on a 0.65 cm^2^ capture area, each with a 55 *µ*m diameter and a spot center-to-center distance of 100 *µ*m. After initial quality control and filtering, 18,045 genes remained, from which we selected the top 5,000 highly variable genes for CCC analysis. Several tumor subtypes were previously detected in [10]. Those were broadly classified into two ductal carcinoma in situ regions (DCIS #1 and #2) and one invasive tumor region (invasive, mixed/invasive) (Fig. 2a).

**Figure 2:**
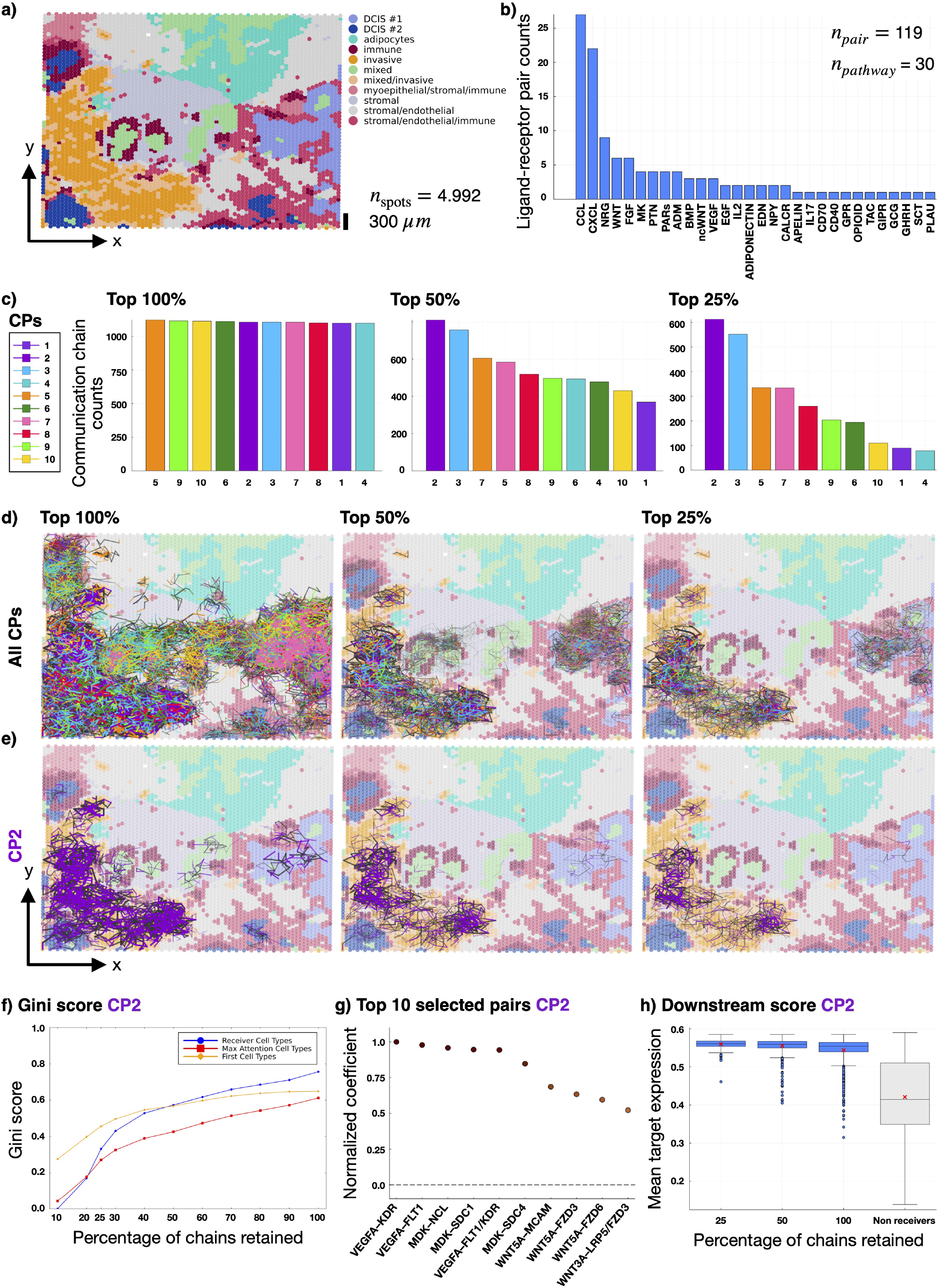
scCChain reveals invasive communication program hotspots in human breast cancer. **a)** Spatial map of Visium CytAssist spots colored by tumor and microenvironmental subtypes. **b)** Ranked counts of feasible ligand–receptor pairs per signaling pathway with total numbers of pairs and pathways indicated. **c)** Communication chain counts per communication program (CP) after transformer-based prioritization for the top 100%, 50% and 25% of chains. **d)** Spatial visualization of all CP-specific chains with line width and transparency scaled by prediction error at different filtering levels (100%, 50% and 25%). **e)** As in d, but for CP 2 only. **f)** Gini scores for CP 2 spot classes as a function of the percentage of retained chains for first, maximum-attention, and receiver spots. **g)** Normalized coefficients of the top 10 ligand–receptor pairs contributing to CP 2, derived from the dimensionality reduction model. **h)** Downstream gene activity scores for CP 2 receiver spots across filtering levels (100%, 50% and 25%) compared with non-receiver spots.

For the CCC analysis, we aimed to spatially localize CP activity associated with secreted ligand–receptor signaling, as defined in CellChatDB, using the set of highly variable genes. To define a candidate communication space, we filtered CellChatDB for feasible ligand–receptor pairs among thes active genes, yielding 119 candidate pairs across 30 signaling pathways (Fig. 2b). Among these pathways, chemokine signaling via the CCL and CXCL families contributed the largest number of candidate pairs for subsequent CP analysis, followed by growth factor pathways including NRG and EGF acting through ERBB receptors, WNT, FGF, VEGF and the heparin-binding growth factors midkine (MK) and pleiotrophin (PTN). These pathways have all been implicated in immune cell recruitment, tumor growth, invasion and angiogenesis [35, 36, 37, 38].

The scCChain workflow initially resulted in 11,083 communication chains for 10 different CPs. For every CP, we sampled five chains with at most five edges from each starting cell, retaining only chains that contained an communication edge. Before communication prioritization via transformer prediction performance, the numbers of communication chains were approximately uniform across CPs (Fig. 2c). Upon prioritization, chains for CP 2 achieved the lowest prediction error among all CCC candidates, closely followed by CP 3, both of which were least affected by the filtering.

To characterize the chains prioritized by transformer performance, we visualized all CP-specific chains in tissue, scaling line width and transparency by prediction error to highlight regions with lower errors (Fig. 2d). Filtering out high-error communications revealed the invasive tumor and DCIS #1 regions as communication hotspots, whereas only a small fraction of chains in the DCIS #2 and immune-spot regions remained among the top 25% lowest-error chains.

Building on CP prioritization, we examined the spatial localization and composition of the signaling mechanisms captured by CP 2. Visualizing CP 2 chains at different filtering levels revealed a strong concentration within parts of the invasive tumor region (Fig. 2e). To quantify this effect, we computed the normalized Gini impurity over the distribution of cell-type classes for the first, maximum-attention, and receiver spots in the chains (Fig. 2f). As filtering becomes more stringent (i.e., fewer chains are retained), the Gini score decreases across all three spot roles, indicating that the retained chains become progressively concentrated within fewer cell-type classes, predominantly the invasive tumor class.

In addition, the model-selected signaling composition of CP 2 was enriched for ligand–receptor pairs with well-established roles in tumor-associated signaling (Fig. 2g). The top-ranked interactions were VEGFA–VEGFR pairs (VEGFA–KDR, VEGFA–FLT1 and VEGFA–FLT1/KDR), consistent with a strong VEGF-driven angiogenic program and the central role of VEGF in tumor neovascularization and progression [37]. The next most prominent group comprised midkine interactions with nucleolin and syndecans (MDK–NCL, MDK–SDC1, MDK–SDC4), in line with reports that midkine, a heparin-binding growth factor, promotes tumor cell survival, invasion, and angiogenesis through these receptors and co-receptors [39]. Finally, several WNT interactions (WNT5A–MCAM, WNT5A–FZD3, WNT5A–FZD6, WNT3A–LRP5/FZD3) were enriched (albeit ranked lower), consistent with the established role of WNT signaling in driving cancer cell migration, invasion, and stemness [36].

Communication chains were generated using distance-aware transcriptional similarity and ligand-receptor coexpression, whereas the communication prioritization model operated on all highly variable genes. This setup allows the model to learn more complex dependency structures along the *a priori* constructed chains that match senders to receivers, for example downstream transcriptional responses in receiver cells after a functional ligand–receptor binding event. To quantify downstream effects, we computed a downstream activity score for each CP based on genes annotated as downstream of the receptor genes in the top three prioritized ligand–receptor pairs, following [21]. For CP 2, the downstream activity score in prioritized receiver spots, calculated from downstream genes of *KDR, FLT1* and *NCL*, increased on average with stronger filtering (Fig. 2h).

Collectively, these results demonstrate that scCChain reveald an invasive, pro-angiogenic CP in a human breast cancer tissue sample, consistent with prior knowledge, and pinpointed the spatial hotspots within the invasive tumor region.

### scCChain supports targeted interaction analysis at single-cell resolution

scCChain is applicable to high-resolution imaging-based spatial transcriptomics, such as Xenium In Situ. Combined with cell segmentation algorithms [40, 41], these platforms yield single-cell–resolved measurements that are well suited for spatial CCC analysis. Because imaging-based spatial transcriptomics platforms typically use targeted panels covering only hundreds to a few thousand pre-selected genes, they offer more limited transcriptome breadth than sequencing-based assays and are therefore less suited to exploratory CP discovery [10, 42]. For such settings, scCChain supports targeted analyses for user-specified ligand–receptor pairs.

We illustrate this targeted analysis using the first replicate of the first sample from a human breast cancer Xenium dataset, generated from an adjacent serial section of the same tumor specimen from the same individual as the Visium dataset analyzed above (Fig. 3a). After preprocessing, the dataset comprised 163,565 cells and 313 profiled genes. Only three secreting ligand–receptor pairs from CellChatDB were detected for this dataset, including CXCL12–CXCR4, a chemokine interaction implicated in immunecell recruitment and tumor proliferation [35]. We focused on this ligand–receptor pair for downstream analyses.

**Figure 3:**
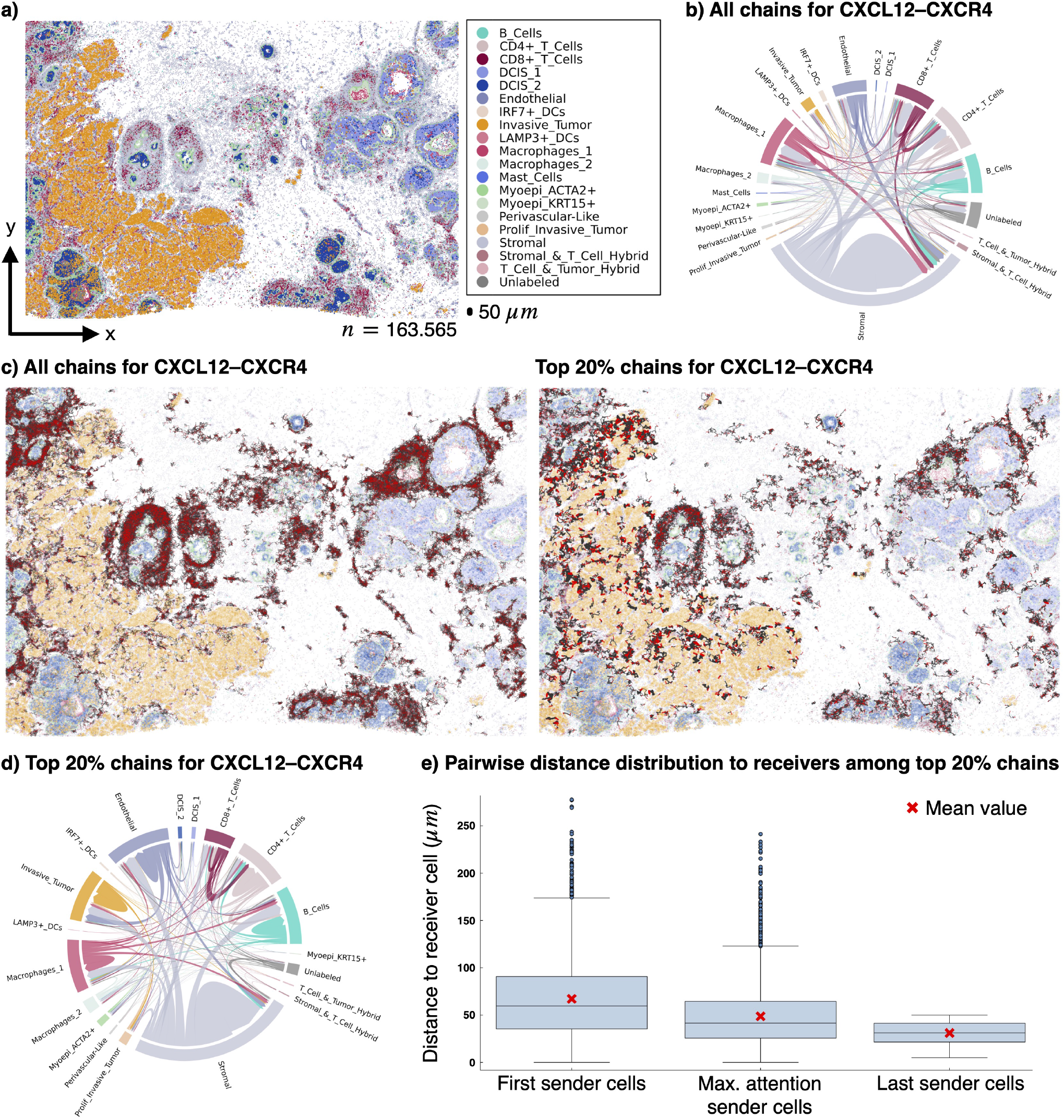
Targeted CXCL12–CXCR4 signaling analysis in Xenium In Situ data. **a)** Human breast cancer Xenium In Situ spatial transcriptomics dataset with cells colored by annotated cell type. The black scale bar below the legend corresponds to 50 *µ*m distance. **b)** Communication chord diagram summarizing all initially sampled CXCL12–CXCR4 communication chains, based on the first sender cell and the receiver cell in each chain. Arrows indicate the direction of candidate communications from sender to receiver cell types. **c)** Spatial plots of CXCL12–CXCR4 chains: left, all generated chains, and right, the 20% of chains with the lowest transformer model prediction error, with line width and opacity scaled inversely with error. **d)** Communication chord diagram for the 20% lowest-error chains based on the most-attended sender cell and the matched receiver cell in each chain. **e)** For the 20% lowest-error chains, pairwise distance distributions between receiver cells and, from left to right, first sender cells, most-attended sender cells, and last sender cells in each chain.

Using a distance-aware cell graph with a 50 *µ*m per-cell coverage radius, we initially sampled 22,068 communication chains with up to 20 cells each (Fig. 3d,e). Incorporating cell-type annotations, we summarized the resulting sender-receiver pairs in a chord diagram (Fig. 3b). Stromal cells were the most frequent senders, consistent with their established role as CXCL12 producers that recruit CXCR4^+^ immune cells, including T and B lymphocytes, in the tumor microenvironment [35]. Mapping all initially generated chains onto the tissue revealed that putative CXCL12-CXCR4 communication was concentrated in tumor-associated regions, with dense sets of chains surrounding both DCIS regions (Fig. 3c, left), suggesting elevated immune activity in these niches.

Transformer-based prioritization of chains reshaped this picture. Although chains in DCIS regions remained more abundant among the 20% of chains with the lowest prediction errors, chains in the invasive tumor region showed the lowest errors overall, as reflected by their thicker lines and higher opacity (Fig. 3c, right). A chord diagram constructed from the most-attended sender cells and matched receiver cells among these prioritized chains showed a relative increase in communication events involving invasive tumor and endothelial cells, including a substantial fraction of tumor-to-tumor communication, and a reduced contribution from stromal and CD8^+^ T-cell communication events compared with all chains (Fig. 3d). This shift is consistent with reports that the CXCL12–CXCR4 axis acts in both paracrine and autocrine loops within the tumor compartment, where CXCL12-producing tumor cells signal to CXCR4^+^ tumor cells to promote growth, invasion, and angiogenesis [35, 43].

We next examined at which spatial ranges the model places the highest weight on CXCL12-CXCR4 communication events. Using aggregated attention weights, we compared the distances from receiver cells to (i) the first sender cell in each chain, (ii) the sender cell with maximum attention, and (iii) the last sender cell (Fig. 3e). Attention tended to concentrate on intermediate-range senders. Most-attended sender cells were, on average, closer to receivers than the first cells in the chains (48.67 *µ*m vs. 67.13 *µ*m), but farther away than the last sender cells (31.05 *µ*m). This pattern suggests that CXCL12-CXCR4 signaling is most informative within a limited spatial range on the order of the chain-construction radius, rather than being dominated by either the most proximal or the most distal senders.

To pinpoint primary target cell populations of CXCL12–CXCR4 signaling and their spatial localization, we further stratified prioritized chains by receiver type (Fig. 4a). Starting from the most frequent receiver type and successively adding the next most abundant types until 80% of prioritized receivers were covered, we obtained six major receiver types: B cells, CD4^+^ T cells, endothelial cells, invasive tumor cells, macrophage 1 cells, and stromal cells. Spatial maps of chains for each receiver type revealed distinct localization patterns (Fig. 4a). Chains with invasive-tumor receivers were concentrated in the invasive region, chains with immune or stromal receivers predominantly surrounded the DCIS regions, and chains with endothelial receivers were scattered across both invasive and DCIS areas.

**Figure 4:**
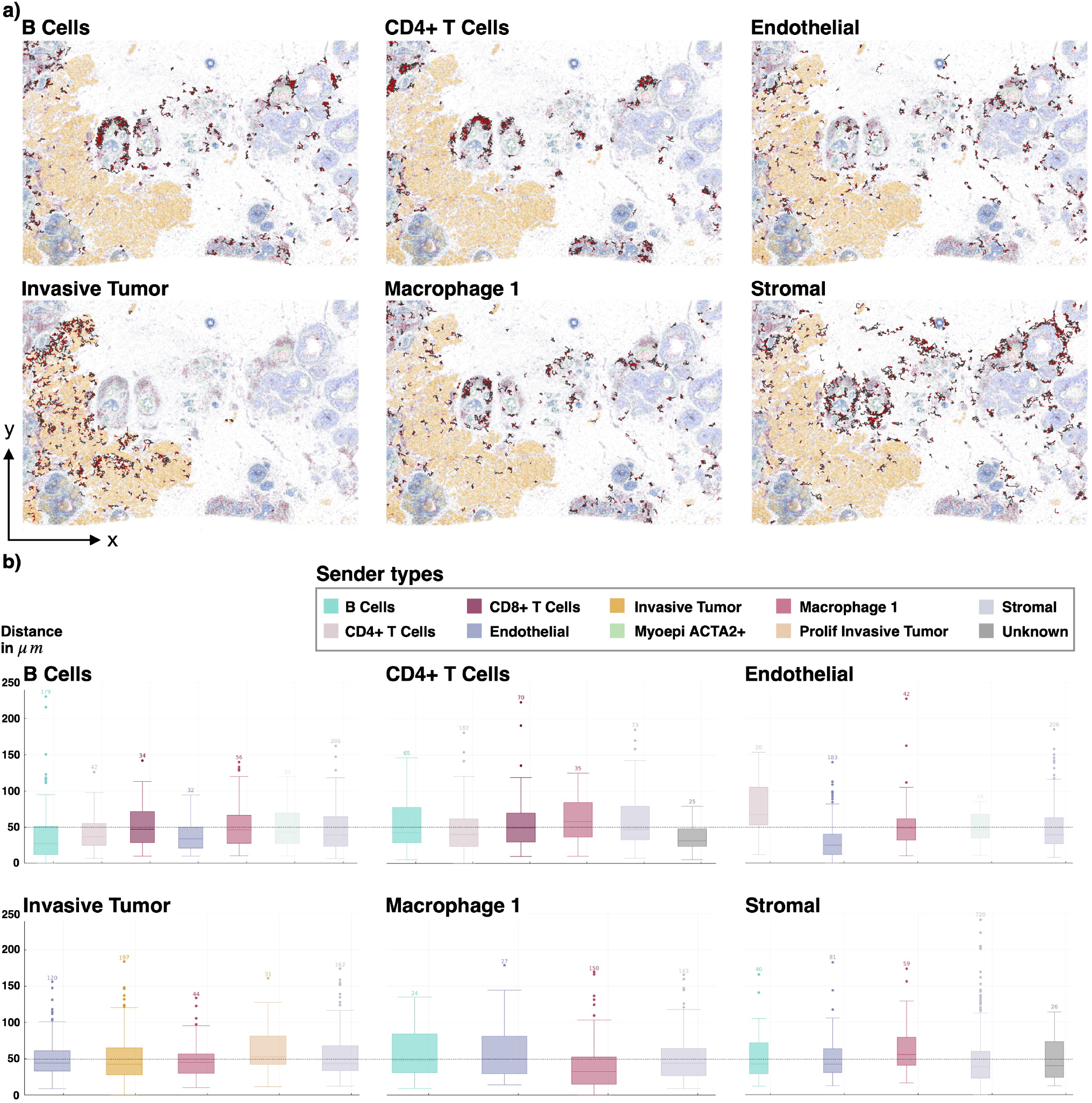
Receiver type–stratified analysis of prioritized communication chains. **a)** For every selected receiver type: Communication chain plot in tissue for all chains where the last cell is of the receiver type. Communication edges are plotted in red, similarity edges in black. **b)** Per receiver cell type: Distribution of pairwise distances between the most attended sender cell and the receiver cell per communication chain. Each box represents the most attended sender cells from a specific sender type. Lower box values reflect closer spatial proximity between model-selected sender and receiver cells. Sender types with less than 20 counts were removed. The black dashed line indicates a distance of 50 *µ*m.

We characterized sender populations and communication distances for each major receiver type by examining the most-attended sender cells in the prioritized chains (Fig. 4b). For every receiver type, we summarized the distance to the receiver cell, stratified by sender cell type. Across receiver types, stromal cells were identified as consistent senders, with median distances just below 50 *µ*m, close to the chain-construction radius. Distance distributions for the most-attended senders were broadly similar across sender and receiver types, with most medians lying below 50 *µ*m, although a minority of chains extended beyond 100–200 *µ*m. Notably, communication events, in which the sender and receiver cell shared the same cell type, showed the shortest median distances for all receiver types except CD4^+^ T cells, for which cells labeled as Unknown were the closest senders.

This analysis demonstrates that scCChain localizes correctly communication niches for selected ligand–receptor pairs at single-cell resolution, to identify predominant sender-receiver cell-type pairs, and to characterize their spatial ranges.

## Discussion

We here introduced scCChain, a computational framework that reformulates spatial CCC as a sequence modeling problem. By building distance-aware communication chains and compressing ligand–receptor pairs into interpretable CPs, scCChain maps candidate interactions to specific tissue locations and ranks chains by their explanatory power for receiver-side signals. Across the datasets analyzed here, the framework supported both exploratory CP discovery and targeted analysis of specific ligand–receptor pairs, while remaining compatible with different spatial transcriptomics platforms.

Conceptually, scCChain combines three components that together address key challenges in computational analyses of spatial CCC. First, communication chains replace both the pair-based view of CCC, in which communication events are modeled independently for sender-receiver or cell-type pairs [14, 15, 18], and fixed-radius or multi-hop spatial graphs [17, 19, 21] with directed paths through tissue. This design allows models to incorporate neighborhood information over variable spatial ranges while keeping each unit compact and computationally tractable for transformer architectures. At the same time, the guilty-by-association principle lets lowly expressing, or communication-silent cells, contribute contextual information. Second, structured dimensionality reduction over ligand–receptor-specific communication layers yields non-negative, sparse, and complementary CPs, where loadings directly correspond to contributions of individual ligand–receptor pairs [26, 27]. Third, transformers trained to predict receiver-cell expression from sender cells exploit broader gene-expression dependency structure beyond genes covered in curated ligand–receptor pair databases, and their attention weights provide a complementary view on which senders are most informative for a given receiver [25]. Together, these elements yield a representation that is biologically interpretable, scalable to large-scale high-resolution spatial transcriptomics datasets, and aligned with modern sequence-modeling architectures.

Analyses across single-cell and spot-based spatial transcriptomics data illustrate how scCChain supports both exploratory and targeted spatial cell-cell communication analysis [10]. In Visium breast cancer, scCChain highlighted an invasion-associated program enriched for angiogenic and pro-migratory cues including VEGF-related signaling [37]. This supports the view that CPs capture coordinated signaling modules rather than isolated ligand–receptor pairs. Prioritized receiver spots also showed increased expression of downstream genes linked to the program’s receptors, which provides indirect support that inferred interactions align with transcriptional consequences of active signaling. In Xenium, targeted analysis of CXCL12–CXCR4 signaling demonstrated that prioritization can refine broad spatial co-localization toward highly informative niches in invasive tumor and endothelial contexts [35, 43]. Across receiver types, stromal cells remained consistent senders, while attention emphasized intermediate distance contributors, suggesting that spatially structured microenvironments can contribute strongly to receiver side signals beyond immediate nearest neighbors.

Some limitations of the current framework are important to acknowledge. As with other computational methods for CCC, scCChain relies on curated ligand–receptor resources and mRNA measurements as proxies for protein abundance and signaling activity, which can introduce incompleteness and bias into the curated ligand–receptor repertoire [7, 16]. For high-resolution imaging-based platforms, cell-level profiles further depend on segmentation and transcript assignment, so segmentation errors can propagate into inferred chains [40, 41]. In this work, we used the provided segmentation without explicit correction. Therfore, inferred communication patterns should be interpreted in light of possible mis-segmentation. More fundamentally, spatial transcriptomics assays provide only indirect evidence of communication events, and there is currently no broadly accepted ground truth for evaluating CCC at scale [7]. We therefore rely on model prediction error and biological plausibility as indirect criteria. With the current transformer architecture and optimization strategy, it is not possible to fully disentangle similarity-driven co-variation from shared niche effects and true communication-driven responses. This limitation may contribute to the patterns observed in the Xenium analysis. Chain construction also involves design choices. Parameters such as neighborhood radius, maximum number of steps, and the transition probability between similarity and communication layers determine the spatial scales and cells that chains can reach, and may require tuning for different tissues, resolutions, and technologies. Furthermore, our present transformer model implementation does not include an additional positional encoding. While this could in principle further bias the model toward local dependencies, we chose not to use them here, motivated by a recent study showing that tissue organization can be inferred from dissociated gene-expression profiles by learning spatial priors from spatial transcriptomics data [33]. Finally, the current implementation is not designed for joint analysis across multiple samples or modalities.

These limitations point to several directions for future work. Incorporating prior knowledge on intracellular signaling pathways, transcriptional targets, and cell-type identities into the transformer modeling could help disentangle similarity-driven co-variation from communication-driven responses in receiver cells. In tumor tissue in particular, candidate interactions are often structured by anatomical subregions such as tumor–stroma interfaces and tumor boundaries, which are not explicitly encoded when chains are built from spatial distance alone. An important extension would therefore be anatomy-aware chain construction that leverages histology-derived segmentation masks or morphological annotations to discourage or forbid chain steps across tissue boundaries, or to refine distance weights along anatomically valid paths. While the current similarity-aware graph can implicitly reduce some boundary violations by favoring transcriptionally coherent transitions, explicit anatomical constraints could further improve specificity and interpretability. Further extending chain construction to allow multiple communication edges per chain would enable the study of higher-order communication patterns, as in recent work on relay networks in spatial CCC [21]. In addition, integrating modalities such as single-cell protein measurements and adapting scCChain to learn jointly across multiple tissues and individuals may improve the specificity of communication prioritization and reveal recurrent CPs.

Within these constraints, scCChain provides a compact and interpretable framework for spatially mapping and prioritizing candidate CPs and targeted ligand–receptor pairs across spatial transcriptomics technologies to driving understanding of how environmental cues such as signaling molecules shape tissue physiology and pathophysiology.

## Methods

### Spot-level spatial transcriptomics data preprocessing

Human breast cancer Visium CytAssist data were obtained from https://www.10xgenomics.com/products/xenium-in-situ/preview-dataset-human-breast and were published in [10]. In addition, previously annotated spot types were also retrieved from https://www.10xgenomics.com/products/xenium-in-situ/preview-dataset-human-breast [10]. The data comprise 4,992 hexagonal spots covering a 6.5 × 6.5 mm (0.42 cm^2^) capture area, with expression quantified for 18,056 genes. Each spot has a diameter of 55 *µ*m and the center-to-center spot-distance is 100 *µ*m. We used scanpy for preprocessing the data [44]. We removed spots with fewer than 300 genes expressed and genes expressed in less than 5 spots. After filtering, 4,982 spots and 18,045 genes remained. We next performed highly-variable gene selection using scanpy’s built-in function with setting flavor=seurat_v3 and kept only the top 5,000 highly-variable genes. The raw count data for each spot were then normalized by total counts over all genes for the corresponding cell. Normalized counts were rescaled by 10,000 followed by a log-transformation with adding a pseudo count of 1 using the natural logarithm. We then performed principal component analysis on the per-feature z-transformed data using singular value decomposition, projecting the standardized data onto the first 30 principal components. This embedding served as input for computing the similarity-layer weights used to construct the cell graph underlying communication chain generation.

### Imaging-based spatial transcriptomics data preprocessing

Human breast cancer Xenium In Situ data were obtained from https://www.10xgenomics.com/products/xenium-in-situ/preview-dataset-human-breast and were published in [10]. In addition, previously annotated cell types were also retrieved from https://www.10xgenomics.com/products/xenium-in-situ/preview-dataset-human-breast [10]. For our analyses, we used the first replicate of the first sample, capturing a tissue area of approximately 0.41 cm^2^. The data were pre-segmented into 167,780 cells with expression measured for 313 genes. We used scanpy for preprocessing the data [44]. Specifically, we removed cells with fewer than 10 genes expressed. After filtering, 163.565 cells remain. The raw count data for each cell were normalized by total counts over all genes for the corresponding cell. Normalized counts were rescaled by 10,000 followed by a log-transformation with adding a pseudo count of 1 using the natural logarithm prior to interaction analysis. Due to the limited gene panel, no further gene selection or dimensionality reduction was applied before communication chain generation.

### Ligand–receptor pair database construction

We use the CellChatDB [14] as the default ligand–receptor pair database for scCChain (https://github.com/jinworks/CellChat/tree/main/data, accessed on October 19, 2025). The database can be filtered to include only user-selected interactions, e.g., interactions corresponding to secreted signaling. To prepare the database for CCC analysis with scCChain, we extract the ligand–receptor pair names, along with the corresponding pathway, ligand, and receptor names (which may be multi-subunit). In principle, alternative and custom databases can also be used for CCC analysis, provided the required information can be extracted. Instructions for preparing and loading custom databases are provided in a tutorial notebook.

### Protein–protein interaction database for downstream signaling scores

The protein–protein interaction data used to compute downstream scores for receptor genes (human_signaling_ppi.csv and mouse_signaling_ppi.csv) was retrieved from CellNEST [21] on GitHub (https://github.com/schwartzlab-methods/CellNEST/tree/main/database, accessed on September 23, 2025). Briefly, the directed protein-protein interactions were originally obtained from the NicheNet database [32] and interactions were queried against STRING (v12.0) [45] to obtain experimental confidence scores, where only interactions with positive experimental scores were retained (for details, see [21], Supplementary Information, Supplementary Note 7).

### Cell graph construction

Let 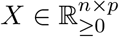 be a gene expression matrix with log-normalized counts, where rows correspond to observations, i.e., cells, spots or bins, and columns to genes. Further, let *P* ∈ ℝ^*n×m*^ denote the matrix consist-ing of the spatial center coordinates of cells (usually *m* = 2). For better interpretation, it is assumed that coordinates are in units of *µ*m. For ease of notation, let *D* = {*𝓁*_1_, …, *𝓁*_*k*_}denote the index set of a curated ligand–receptor pair database possibly consisting of hundreds up to thousands of ligand–receptor pairs. Each *𝓁* ∈ *D* specifies a (possibly multi-subunit) nonempty ligand gene set *A*_*𝓁*_ ⊂ {1, …, *p*} and a (possibly multi-subunit) nonempty receptor gene set *B*_*𝓁*_ ⊂ {1, …, *p*}.

Communication chains are sampled by a random walk approach from a weighted multi-layer cell graph. To build this graph, let *V* = {*v*_1_, …, *v*_*n*_}be the set of nodes (representing cells, spots, or bins) and let *L* = {sim} ∪ *D* be the layer index set, where sim denotes the similarity layer and each *𝓁* ∈ *D* denotes a directed communication layer for a ligand–receptor pair. Edges in each layer are restricted to spatial neighbors defined from *P*, and edge weights decay with Euclidean distance between nodes. The edge set is

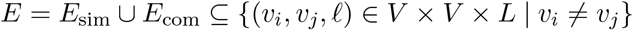

which by definition excludes s elf-loops. The weighted cell graph is 𝒢 = (*V, L, E, w*) where *w*: *E*→ ℝ denotes the edge weight function for the different layers.

To scale the edge weights by spatial distance scores of node pairs, we define

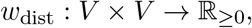

by a truncated Gaussian kernel with bandwidth parameter 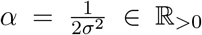 and maximum distanceparameter *d*_max_ ∈ ℝ _*>*0_

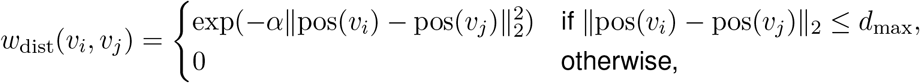

where pos: *V* → ℝ ^*m*^ maps every node to its spatial position pos(*v*_*i*_) = *P*_*i*_. The bandwidth parameter controls the decay behavior of the kernel function with increasing distance of nodes. Increasing the parameter results in faster decay. Truncating the kernel at *d*_max_ sparsifies the cell graph and increases computational efficiency by avoiding computation and storage of edge layer weights for all cell pairs, i.e.,

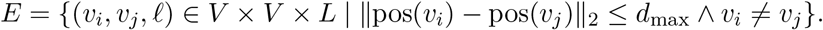

To define the distance-aware edge weight function for the similarity layer, let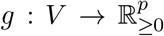 map each node *v*_*i*_ ∈ *V* to its corresponding gene expression vector *g*(*v*_*i*_) = *X*_*i*·_. Let *f*: ℝ ^*p*^ → ℝ ^*q*^, with *q* ≪ *p*, be a previously fitted embedding function derived for example from the encoder of an autoencoder-based model like scVI [46], or from the projection obtained by principal component analysis. We define the embedded representation of a node *v*_*i*_ as

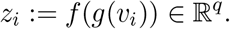

Further, we denote by cos_sim the cosine similarity between two embedded node representations, defined as

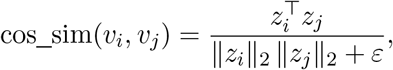

where *ε* = 10^*−*12^ is a small positive constant for numerical stability. The distance-aware edge weight function for the symmetric similarity layer is then defined by

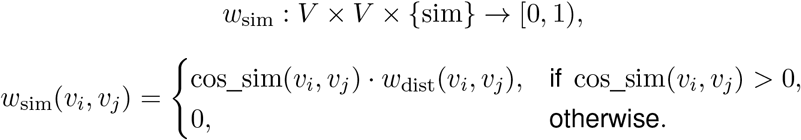

Truncating to positive cosine similarity values ensures nonnegative weights, which is required when subsequently transforming edge weights into transition probabilities for random-walk sampling.

Next, for a node *v*_*i*_ ∈ *V* with log-normalized expression vector 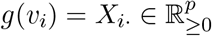, we define the subunit-aware geometric-mean expression of a nonempty gene set *G* ⊆ {1, …, *p*} by

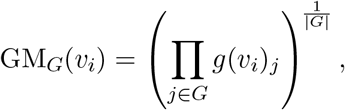

ensuring all subunits have to be expressed for a nonzero score. The distance-aware, directed communication weights are obtained by the function

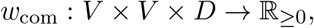

With

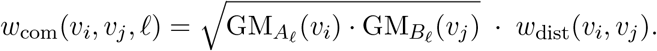

The edge weight function across layers is defined by

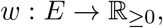

With

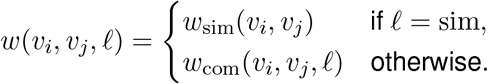

### Communication program discovery

To identify CPs, we perform structured dimensionality reduction on the communication layer edge weights from the cell graph 𝒢, aggregating the possibly extensive number of |*D*| ligand–receptor pairs in a reasonable number of CPs. The objective is to learn a low-dimensional latent representation of cell pair communication scores that satisfies three core criteria:

a. Disentanglement: Each latent dimension captures distinct, complementary communication information.
b. Interpretability: Each CP is characterized by a sparse subset of ligand–receptor pairs, each associated with a coefficient (loading) quantifying its contribution.
c. Positive support: The representation scores should be interpretable as communication scores for the cell pairs.

To obtain such a representation, we use the Boosting Autoencoder (BAE), including a disentanglement constraint for the latent dimensions [26], that has been previously adapted to model CCC data [27]. Briefly, the BAE integrates componentwise boosting into parameter optimization of an autoencoder model for updating the weights of a single-layer neural network encoder. All other model parameters are jointly optimized via stochastic gradient-based optimization. Let *h* denote the number of latent dimensions of the BAE encoder. During optimization and starting from a zero initialization of the encoder weight matrix *W*_sparse_ ∈ ℝ^*h×*|*D*|^, the BAE selects features in a stepwise manner over training epochs, resulting in a sparse mapping from the ligand–receptor space to the *h*-dimensional latent space while enforcing disentanglement of latent dimensions

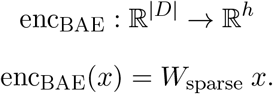

To further enhance interpretation and to enable automatized soft-clustering, the encoder can be extended by a split-softmax function [27]

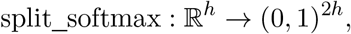

With

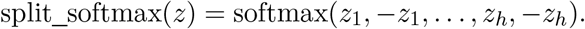

CP candidates are defined by the 2*h* output dimensions of the extended BAE encoder. Representations of cell pairs can be assigned to CPs by determining the arg max-value among all 2*h* dimensions. Lig- and–receptor pairs are directly linked to CPs by the sparse encoder weight matrix *W*_sparse_, which can be interpreted as loadings.

Importantly, being optimized in a fully data-driven manner, not all candidate CPs necessarily capture biologically relevant signals. To filter redundant CPs, we apply a criterion based on Youden’s *J* statistic (*J* = TPR − FPR) using as ground-truth labels whether each cell pair is assigned to the CP or not [47]. Specifically, we retain a CP if its *J* -maximizing threshold achieves sufficiently strong separability between cell pairs assigned to that CP and all others, as determined by a global threshold on the maximum *J* scores. In addition, each CP must satisfy a minimum support, that is, a specified minimum number of assigned cell pairs, to be retained. We denote the set of retained CPs by 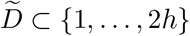.

Let 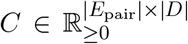 denote the matrix of communication weights, where *E*_pair_ ⊆ *V* × *V* is the set of ordered cell pairs with at least one non-zero interaction. For every pair (*i, j*) ∈ *E*_pair_ and ligand–receptor pair-index *𝓁* ∈ *D*, we define

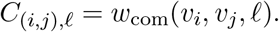

Applying the BAE encoder row-wise to *C* yields, for each (*i, j*) ∈ *E*_pair_,

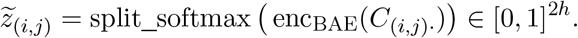

For 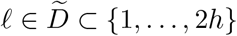, the CP-specific edge weights are defined by

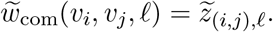

The retained CPs induce a summarized communication graph that we use for communication chain generation. Let

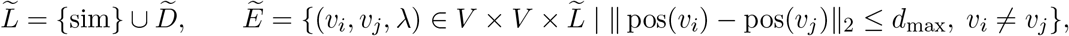

and define

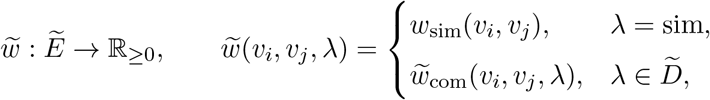

resulting in the summarized cell graph

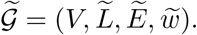

### Communication chain generation

Communication chains are generated by a weighted random walk either (i) from the summarized graph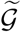 for CP analyses, or (ii) directly from the original graph 𝒢 for targeted communication analyses. Because the procedure is identical, we use the generic notation 𝒢.

For a feasible communication mechanism *𝓁* with layer set *L*^(*𝓁*)^ = {sim, *𝓁*}, let 𝒢^(*𝓁*)^ = (*V, L*^(*𝓁*)^, *E*^(*𝓁*)^, *w*^(*𝓁*)^), where 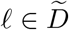for CP analysis and *𝓁* ∈ *D* for targeted analysis with edge set

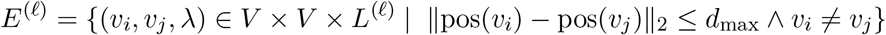

and weight function

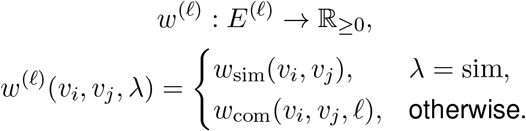

The chain generation process starts from nodes with high outgoing signaling potential. Therefore, let

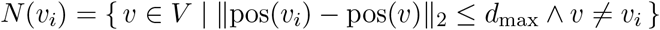

be the neighborhood of a node *v*_*i*_ and let

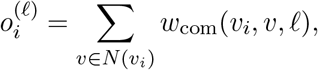

define the outgoing communication potential from *v*_*i*_ to its neighbors. We define the set of feasible starting nodes by

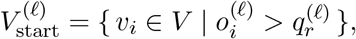

where 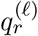 is the *r*-quantile of the nonzero scores

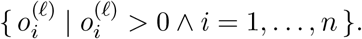

Next, let (*X*_*t*_)_*t*≥1_ define the weighted random walk for generating communication chains and let (*Z*_*t*_)_*t*≥2_ be i.i.d. Bernoulli(*ρ*) with *ρ* ∈ [0, 1] and *Z*_*t*_ ∈ {0, 1}, independent of (*X*_*t*_)_*t*≥1_. *Z*_*t*_ is then mapped to a process that represents the coin-flip mechanism used to select the layer for selecting the next node

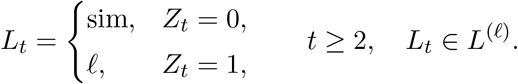

To enforce spatial information accumulation, we set the first layer choice deterministically to the similarity layer,

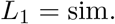

For *λ* ∈ {sim, *𝓁*} we define the random walk transition matrix *P* ^(*λ*)^ on *V* by

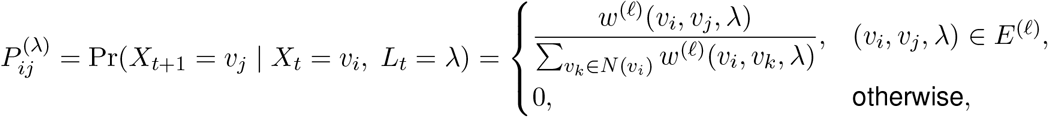

with self-loops excluded by the edge set 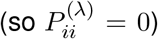. Next, we define a stopping criterion that terminates chain generation either upon the first communication step or once a prespecified maximum number of nodes has been selected. Let *t*_max_ ∈ ℕ_≥2_ denote a maximum number of transitions from one node to another and

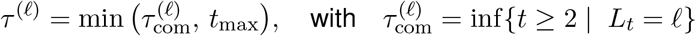

a stopping time for the process (*X*_*t*_)_*t*≥1_. Starting from a node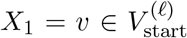, we generate a chain of nodes by

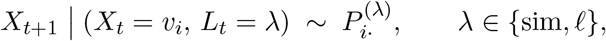

for *t* = 1, 2, … with *t* ≤ *τ*^*(𝓁)*^. The resulting communication chain is the stopped path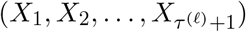. By construction, at most one communication edge can occur, and the procedure halts immediately after its first occurrence or when *t*_max_ is reached.

Communication chains are generated for every feasible starting node in 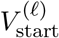 and every feasible communication mechanism *𝓁*. Subsequently, chains without having traversed an communication edge (i.e.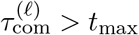) are removed prior to communication chain prioritization.

### Communication prioritization model

The communication prioritization model is based on a transformer architecture [25] and operates on communication chains, where tokens correspond to cells. Its self-supervised objective is to predict the gene expression of receiver cells by identifying which sender-cell gene expression patterns are most informative for the receiver, thereby incentivizing the model to learn cross-cellular gene dependencies relevant to intercellular signaling.

For ease of notation, the forward pass is exemplified on an arbitrary communication chain (*v*_1_, …, *v*_*k*_) of length *k* ≤ *t*_max_ + 1, but note that the model processes multiple communication chains in parallel. We first map cells in the communication chain to their corresponding gene expression vectors

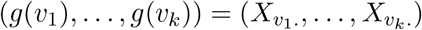

We then split the chain into its sender part, consisting of all cells before the communication step, 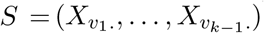 and the receiver part consisting of only the last cell 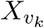. To ensure consistency across all generated sequences, zero vectors are added to the sender part where the number of cells is smaller than the maximum number of transitions *t*_max_ in chains

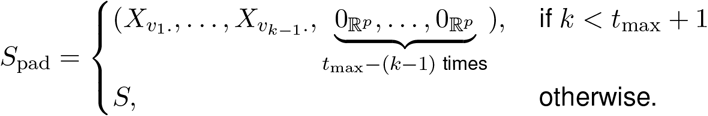

Since cells are already represented by continuous gene expression vectors, we operate directly in gene expression feature space. This differs from natural-language settings, where a finite set of discrete tokens is first mapped to trainable embedding vectors in an arbitrary-dimensional feature space.

The communication prioritization model consists of an autoencoder architecture, reducing the dimensionality of the gene feature space to mitigate noise effects. The encoder consists of a single multi-head scaled dot-product cross-attention layer.

Let *n*_heads_ ∈ ℕdenote the number of heads and let *d*_QK_, *d*_V_ ∈ ℕ with *d*_QK_, *d*_*V*_ ≪ *p* denote the dimensionalities of the queries, keys, and values. For computing the output for a single head *i* ∈{1, …, *n*_heads_} of the cross-attention layer, we define the queries based on the receiver cell, and keys and values are based on the sender cells

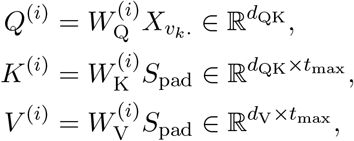

with trainable weight matrices 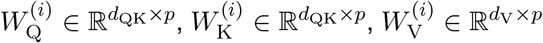.

The attention scores are computed by

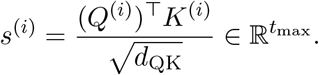

The scores are used to obtain the attention weights *a*^(*i*)^ for building weighted averages of the value representations. Attention weights are constructed by applying a masked softmax to the attention scores, excluding positions corresponding to padded zero vectors. Consequently, padded sender positions are assigned zero weight and do not contribute to the encoded representation. Let 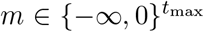 be an additive attention mask with *m*_*j*_ = 0 for valid sender positions and *m*_*j*_ = − ∞ for padded positions. The masked attention scores and weights for head *i* are then given by

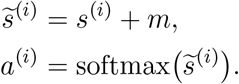

To prevent overfitting and to encourage distributing attention across senders, we apply dropout to the attention weights. Specifically, let *p*_drop_ ∈ [0, 1) denote the dropout rate and let 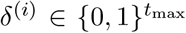be an elementwise dropout mask with 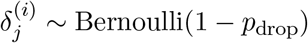. The dropped-out attention weights are

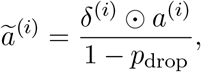

Using these refined attention weights, the head output is computed as a weighted sum of the value vectors

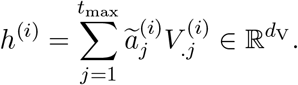

Because the receiver cell provides a single query vector that attends over the sender positions, the attention computation scales linearly in the sender sequence length, 𝒪(*t*_max_), rather than quadratically as in standard attention.

The outputs of individual heads are then concatenated and aggregated by multiplication with a trainable weight matrix 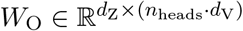to compute the final encoder representation

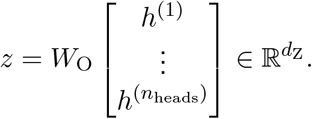

The decoder is a fully connected multilayer feedforward neural network that maps *z* to the predicted gene expression of the receiver cell

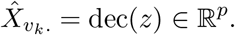

During training, the model is optimized by minimizing the mean squared error given the original receiver cell’s gene expression and the model prediction

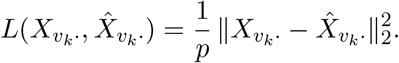

### Prioritization of candidate communication events via model prediction error

Let *n*_chains_ denote the number of communication chains and let 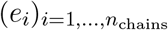 denote the prediction errors produced by the trained communication prioritization model for the chains 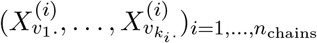. Next, let

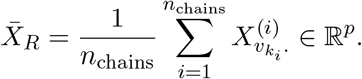

denote the average gene expression vector across receiver cells in the chains and let

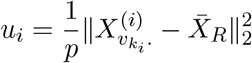

denote the baseline mean squared deviation of its receiver expression profile for chain *i*. We then define the adjusted prediction errors by

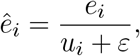

where *ε >* 0 is a small constant added to the denominator to prevent division by zero.

By construction, 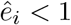 indicates that the communication prioritization model predicts chain *i* better than the baseline mean predictor built from average receiver gene expression. A value of 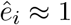 corresponds to baseline-level performance, and 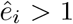 indicates worse-than-baseline performance. We use 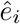 to prioritize communication chains, assigning higher priority to chains with smaller adjusted prediction errors.

### Identification of most-attended sender cells

Given the refined attention weights 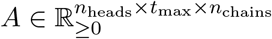 for all communication chains from the trained communication prioritization model, we first averaged attention across heads to obtain 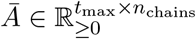

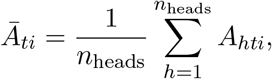

where *t*_max_ is the maximum number of sender cells in a chain. For each chain *i*, the most-attended sender index was defined as

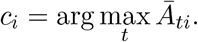

### Error-based weighting of communication chains for visualization

To highlight prioritized communication events in plots where communication chains are overlaid on the tissue, we scaled the width and opacity of lines connecting cells inversely with the corresponding model prediction errors, such that lower-error interactions appear thicker and more opaque. Given a collection of non-negative prediction errors (*e*_*i*_)_*i*=1,…,*n*_ corresponding to communication chains, we first convert them into inverse normalized scores in [0, 1]:

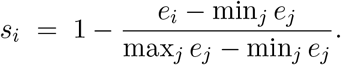

We next bin the scores into *n*_bin_ equally spaced intervals on [0, 1] and assign each chain to one of these bins. The bin index for chain *i* is defined as

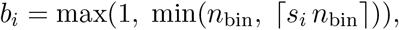

ensuring that all chains fall into one of the *n*_bin_ bins. For each bin *b*, we compute a corresponding line width *w*_*b*_ and opacity *α*_*b*_ using monotonic functions:

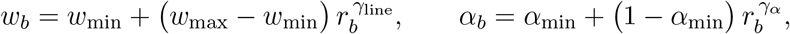

Where

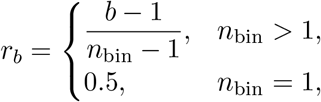

and *w*_min_, *w*_max_, *α*_min_, *γ*_line_, *γ*_*α*_ are user-defined parameters controlling the dynamic range and nonlinear scaling of the visual weighting. Lower-error chains (larger *s*_*i*_) are assigned to higher-index bins and therefore appear thicker and more opaque in the tissue plots, making high-confidence communication events visually prominent, whereas higher-error chains become thinner and more transparent.

### Gini score

We quantify cell type label heterogeneity using a normalized Gini impurity [48]. For categorical labels *m*_1_, …, *m*_*n*_ with optional nonnegative weights *w*_1_, …, *w*_*n*_ (default *w*_*i*_ = 1), the (weighted) class proportions are

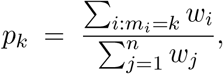

and the standard Gini impurity is

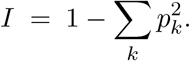

Let *K*_obs_ be the number of observed classes (*p*_*k*_ *>* 0) and *K*_norm_ the number of classes used for normalization, with *K*_norm_ = *K*_obs_ by default or *K*_norm_ = *K* ≥ *K*_obs_ ≥1 if a total of *K* possible classes is specified. We then define the normalized impurity

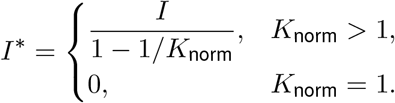

which equals 0 for perfectly pure labels and 1 for maximally heterogeneous labels given *K*_norm_ classes.

### Downstream gene scoring

To quantify downstream activity of receptors, we leveraged the CellNEST intracellular signaling database and applied its downstream-gene scoring strategy [21]. We first compiled species-specific protein–protein interaction networks from the database and extracted all edges annotated with an experimental interaction score. For a given set of source genes (e.g. receptors), first-order downstream genes were defined as the direct targets of these sources in the protein–protein interaction, and second-order downstream genes as the targets of the first-order genes. For each second-order target gene, we aggregated multiple incoming edges by taking the maximum experimental interaction score across all upstream sources, and ranked genes in descending order of this score. We then selected the top 20% of genes, including the immediate first-order targets in the final downstream gene set. Let *X* denote the cell-by-gene (log-normalized) gene expression matrix and *G* the subset of selected downstream genes present in the columns of *X*. For each cell, we computed a downstream activity score as the arithmetic mean of the expression values of genes in *G*.

### Communication prioritization model training

For training the model, we split the data into training (80%) and validation (20%) sets. At the beginning of training, model parameters are randomly initialized [49]. In each training iteration, we sample a mini-batch by drawing a random subset from the training set. Parameter updates are performed using the AdamW optimizer [50]. Training is stopped either when a maximum number of iterations is reached or when the validation loss does not improve for a predefined number of epochs (early stopping). In the latter case, the final model is chosen as the one with the minimal validation loss.

### Statistics and reproducibility

This study is primarily computational and descriptive. CPs, communication chains, and spatial communication hotspots were identified and prioritized using model-based procedures implemented in scCChain rather than formal hypothesis-testing frameworks. For communication program discovery, the boosting autoencoder was trained directly on the standardized cell-pair communication matrix without a train/validation split. Decoder parameters were optimized using AdamW [50], whereas encoder weights were updated by componentwise boosting. Training was performed for a maximum number of epochs and stopped earlier when the change in batch loss fell below a predefined tolerance. For communication prioritization, data were split into training and validation sets, model parameters were optimized using AdamW [50], and the final model was selected according to the minimum validation loss with early stopping.

scCChain is implemented in Julia [51], with Python-based utilities for preprocessing spatial transcriptomics data [44]. The public GitHub repository contains the full source code, environment specifications for Julia and Python dependencies, tutorial notebooks, bundled ligand–receptor databases, figure-reproduction scripts, and detailed instructions for reproducing the analyses presented in this manuscript.

## Data availability

All datasets used in our analyses can be accessed at https://www.10xgenomics.com/products/xenium-insitu/preview-dataset-human-breast. The data were published in [10].

## Code availability

An open-source implementation of scCChain, together with tutorial notebooks on usage and data preparation, is available at https://github.com/NiklasBrunn/ScCChain.jl.

## Acknowledgements

This work was supported by the Deutsche Forschungsgemeinschaft (DFG, German Research Foundation) under Project-ID 322977937 – GRK 2344 (C.L.F., H.B., N.B., T.V.) and Project-ID 499552394 – SFB 1597 (H.B., L.C.G., M.H., N.B.).

## Author contributions

H.B. supervised the project and, together with N.B., conceived the work. N.B. implemented the modeling framework. L.C.G. implemented the chain generation framework with contributions from N.B.. J.K. and

K.F. contributed to the conceptual design and implementation of the interaction prioritization model. N.B. performed the computational analyses. N.B. wrote the manuscript, with contributions from H.B. and L.C.G.. C.L.F., J.K., K.F., L.C.G., M.H., and T.V. critically reviewed the manuscript. All authors reviewed and approved the final version of the manuscript.

## Competing interests

The authors declare no competing interests.

## Supplementary material

**Supplementary Fig. S1:**
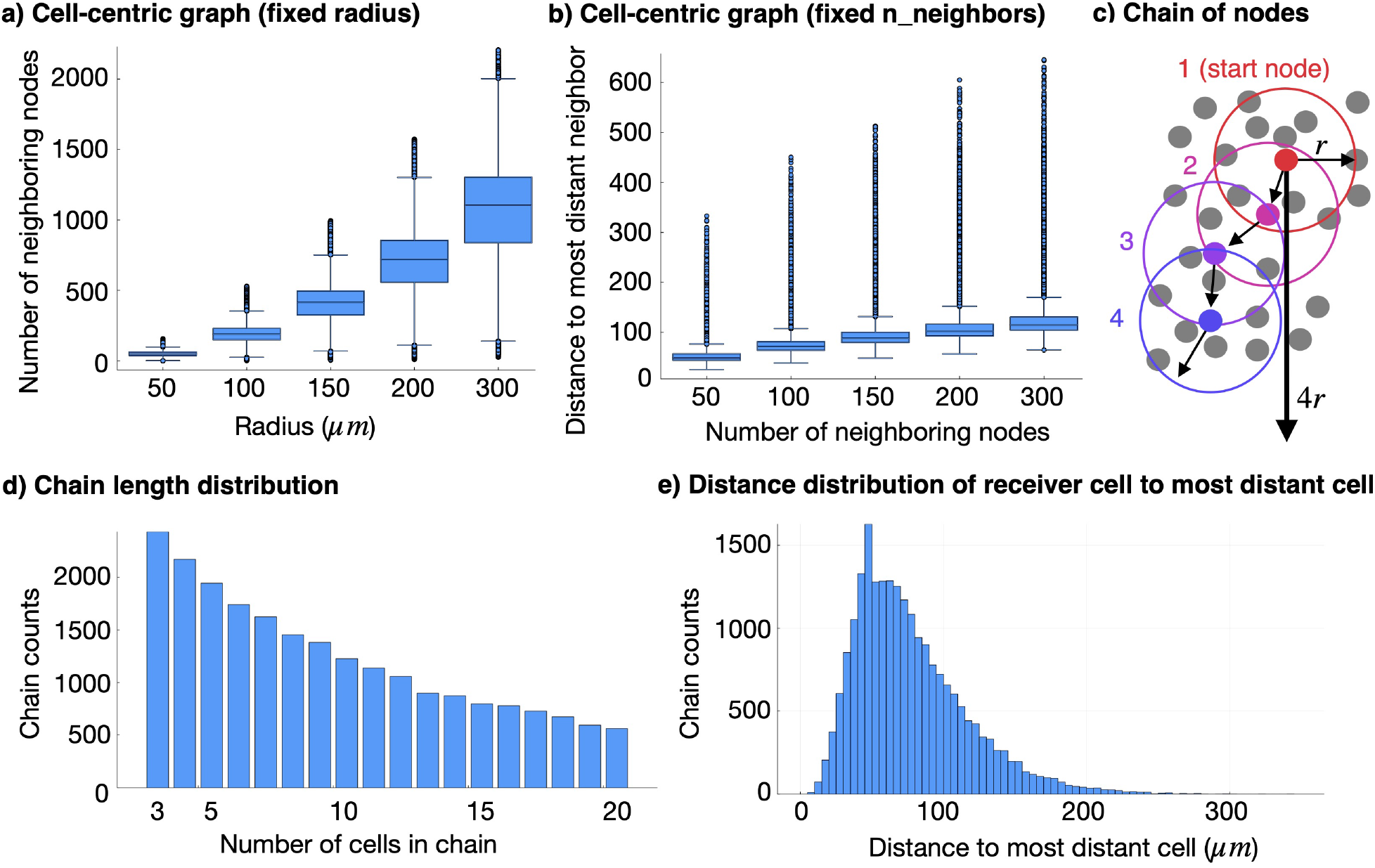
Communication chains vs. node-centric graphs. Illustrative results on Xenium In Situ human breast cancer data. **a)** Distribution of neighbors per cell in a node-centric neighborhood graph across neighborhood radii (*µ*m). **b)** Distribution of distances from the center cell to its farthest neighbor (*µ*m). **c)** Schematic of node-centric shifts used to generate interaction chains of length 4. **d)** Distribution of chain cell counts for one configuration (*r* = 50 *µ*m, *p*_com_ = 0.1, *t*_max_ = 19). **e)** Distribution of distances from the terminal cell (receiver) to the farthest cell (sender) in the chain (*µ*m).

## Notes

### Competing Interest Statement

The authors have declared no competing interest.

